# Convergent mutations in phage virion assembly proteins enable evasion of Type I CBASS immunity

**DOI:** 10.1101/2023.05.21.541620

**Authors:** Desmond Richmond-Buccola, Samuel J. Hobbs, Jasmine M. Garcia, Hunter Toyoda, Jingjing Gao, Sichen Shao, Amy S. Y. Lee, Philip J. Kranzusch

## Abstract

CBASS is an anti-phage defense system that protects bacteria from phage infection and is evolutionarily related to human cGAS-STING immunity. cGAS-STING signaling is initiated by viral DNA but the stage of phage replication which activates bacterial CBASS remains unclear. Here we define the specificity of Type I CBASS immunity using a comprehensive analysis of 975 operon-phage pairings and show that Type I CBASS operons composed of distinct CD-NTases, and Cap effectors exhibit striking patterns of defense against dsDNA phages across five diverse viral families. We demonstrate that escaper phages evade CBASS immunity by acquiring mutations in structural genes encoding the prohead protease, capsid, and tail fiber proteins. Acquired CBASS resistance is highly operon-specific and typically does not affect overall fitness. However, we observe that some resistance mutations drastically alter phage infection kinetics. Our results define late-stage virus assembly as a critical determinant of CBASS immune activation and evasion by phages.

## INTRODUCTION

Bacteria encode a diverse array of defense systems that protect cells from phage infection. One of the most common forms of anti-phage defense is cyclic oligonucleotide-based signaling system (CBASS) immunity, occurring across major phyla and in >16% of all sequenced bacterial genomes^1–3^. In CBASS immunity, phage infection is sensed by a cGAS/DncV-like Nucleotidyltransferase (CD-NTase) enzyme which synthesizes a cyclic nucleotide second messenger signal to control downstream receptors named CD-NTase-associated proteins (Cap) effectors^2,4^. Cap effector activation induces bacterial growth stasis or cell death which inhibits phage replication and thereby prevents propagation throughout the bacterial population^1,4,5^. Bacterial CD-NTase enzymes and Cap effectors are structurally and functionally homologous to human proteins in the cGAS-STING antiviral immune pathway, demonstrating a direct evolutionary connection between core components of human immunity and CBASS immunity in bacteria^2,6,7^

Bacterial CBASS defense systems are extremely diverse in sequence and operon organization. CD-NTase enzymes each share a conserved active site and overall enzyme structure but form eight distinct protein clades A–H which exhibit <10% sequence identity at the amino acid level across CD-NTase clades^2^. The most minimal CBASS operons are comprised of only two genes encoding the CD-NTase enzyme to sense infection and initiate cyclic nucleotide signaling and a Cap effector to recognize the nucleotide signal and trigger abortive infection to prevent phage propagation^1,2,4^. These operons are designated “Type I” CBASS and are distinct from larger Type II, III, and IV CBASS systems which encode additional ancillary proteins proposed to further regulate cyclic nucleotide signaling and immune activation^8^. Previous biochemical and genetic analyses of bacterial CBASS immunity have detailed many important features of cyclic nucleotide-based antiviral signaling and immune evasion by phages^1,2,5,6,9–16^ and have enabled more recent discoveries of a diversity of cGAS-like immune signaling enzymes in vertebrate and invertebrate species^17–19^. However, many studies on bacterial CBASS have relied predominantly on *in vitro* experiments for large-scale analysis of diverse enzymes, or focused experiments on a few, often unrelated Type II and Type III CBASS operons.

A major open question in bacterial CBASS immunity is how CD-NTase enzymes recognize phage infection and initiate antiviral immune signaling. Studies on other anti-phage defense systems have identified specific infection-derived triggers^20–24^ responsible for immune activation, but no detailed mechanism has emerged for initiation of CBASS defense. Many bacterial CD-NTase enzymes are constitutively active *in vitro*^2^, presenting a challenge to understanding how CD-NTases initiate signaling *in vivo*. Diverse models of CBASS activation include recognition of peptides^5^, sensing of non-coding RNAs^25^, conjugation of CD-NTase enzymes to unknown target molecules^15,16^, and indirect detection of metabolite depletion^26–28^.

Here we perform a comprehensive analysis of Type I CBASS immune specificity and evasion by phages. We bioinformatically analyze Type I CBASS and functionally characterize a diverse panel of 15 CBASS defense operons in *E. coli* to demonstrate these compact systems potently restrict dsDNA coliphage replication *in vivo*. Crossing all 15 Type I CBASS operons against 65 diverse coliphages, we create a defense profile network of 975 CBASS defense operon-phage pairings that reveals striking patterns of Type I CBASS-mediated phage restriction. We show that escaper phages acquire resistance to Type I CBASS via spontaneous mutations in the prohead protease, major capsid, and tail fiber virion assembly proteins and that resistance phenotypes are largely operon-specific. We then characterize escaper phage phenotypes and uncover that some structural mutations impact infection kinetics. Together, our results define how phages can alter late-stage viral assembly to evade CBASS immunity and provide a new foundation to explain CBASS specificity and immune evasion by phages.

## RESULTS

### Diverse Type I CBASS operons protect bacteria from phage infection *in vivo*

To characterize the core determinants of CBASS immune specificity, we focused our analysis on Type I CBASS operons devoid of additional signature Cap proteins associated with more complex CBASS operons^8^. We built on previous analysis of CD-NTase sequence diversity in CBASS immunity^2,8,29^ and bioinformatically analyzed 2,376 Type I CBASS operon sequences present in available sequenced bacterial genomes (Figure 1A). Type I CBASS operons are exceptionally diverse and occur in nearly every major bacterial phylum^8^. We created a phylogenetic tree of Type I CBASS based on CD-NTase protein sequence and observed that Type I operons contain representatives from seven out of eight major CD-NTase cyclase clades A– G and include a diverse set of at least 8 distinct Cap effector proteins (Figure 1A). Type I CBASS operons most frequently encode CD-NTase enzymes from clades D, E, and F (>80% of Type I CBASS operons) accompanied by Cap effectors with 2 or 4 predicted transmembrane segments^10^. We observed closely related Type I CBASS CD-NTase proteins encoded adjacent to diverse Cap effectors supporting previous models where exchange of effector proteins facilities rapid evolution of CBASS defense^30,31^.

**Figure 1.**
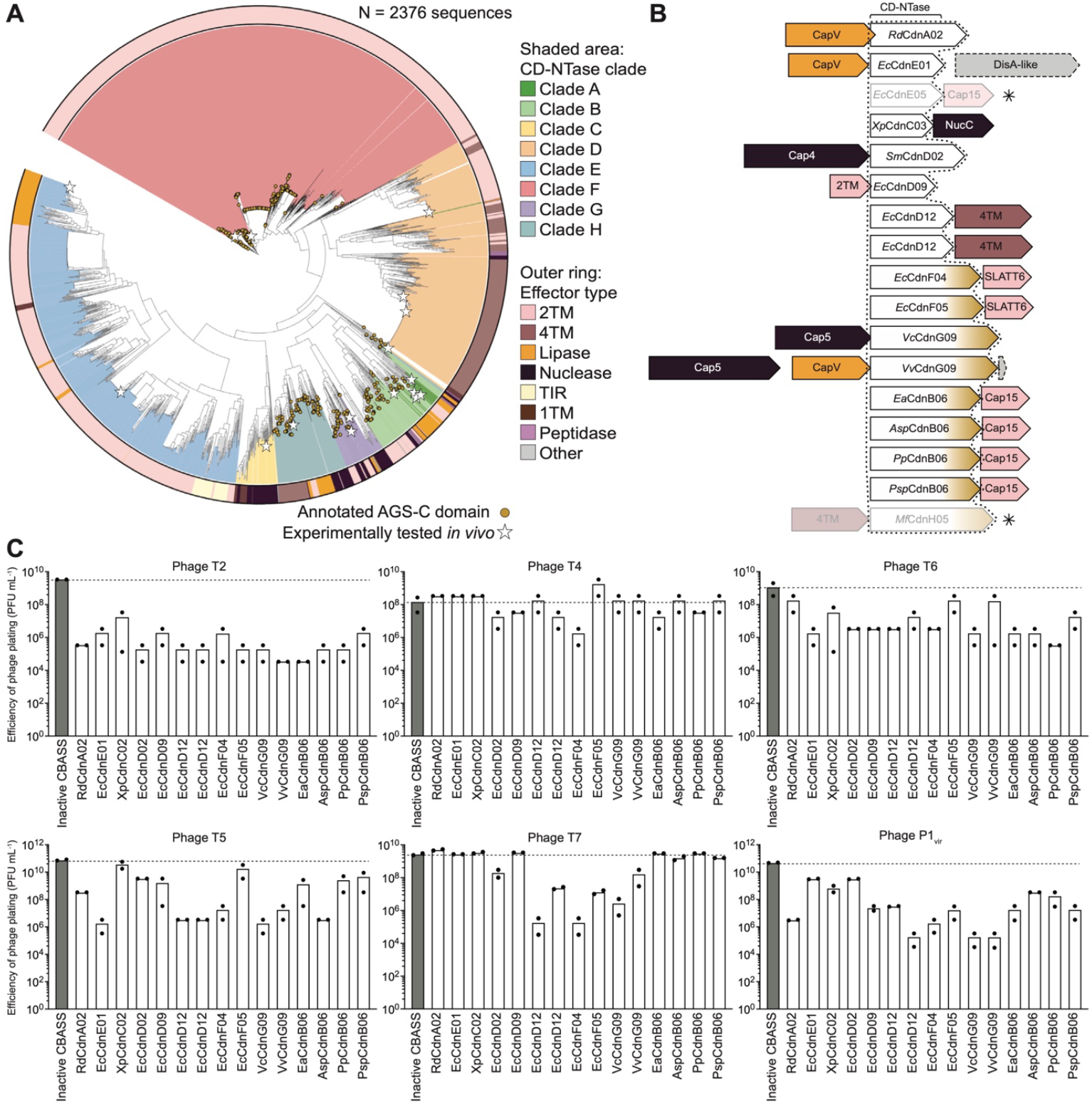
Diverse Type I CBASS operons restrict dsDNA phage replication in vivo. (A) Phylogenetic analysis of ∼2,400 Type I CBASS CD-NTase enzyme sequences. Gold circles denote CD-NTases which are bioinformatically predicted to encode Ig-like AGS-C fusion domains. Stars represent full CBASS operons which were cloned and tested experimentally. (B) Genomic architecture of selected Type I CBASS operons used for further analysis. Cap effectors are labeled based on homology to characterized Cap proteins and colored to match designations in panel (A). Asterisks represent operons which were toxic in *E. coli* and not pursued for further study. (C) Efficiency of phage plating (Y-axis) of six classic lytic or temperate phages on *E. coli* BW25113 cells transformed with the WT CBASS operon indicated on the X-axis or an effector-null CBASS *Ea*CdnB–Cap15ΔTM control^10^.

A distinct feature of Type I CBASS operons is frequent CD-NTase fusion to an ∼18 kDa C-terminal domain of unknown function termed an AGS-C domain^32^. AGS-C domains are predicted to share structural homology with immunoglobulin (Ig)-like fold proteins including vertebrate antibody and nanobody antigen recognition domains^33^ and are a CD-NTase feature not found in other CBASS types. We observed that ∼40% of Type I CBASS operons are predicted to encode a CD-NTase–AGS-C fusion (Figure 1A). Clade B and H CD-Ntases in Type I CBASS almost ubiquitously encode AGS-C fusion domains and there are no examples of this domain in clade A, C, D, and E Type I CD-NTases, whereas clades F and G of the Type I CBASS CD-NTase tree display distinct branches containing or lacking this domain (Figure 1A). Fusion of the AGS-C nanobody-like domain to CD-NTase enzymes suggests a potential role for this domain in modifying the initial steps of viral recognition in CBASS defense.

To begin to define patterns of Type I CBASS defense, we cloned a panel of 17 diverse Type I CBASS operons from gammaproteobacterial genomes and screened each system for the ability to inhibit phage replication in *E. coli*. Our panel of Type I CBASS operons includes CD-NTase representatives from each major clade A–H, examples of operons with and without CD-NTase–AGS-C fusions, and 8 distinct classes of Cap effector proteins (Figure 1B; Table S1). Expression of 15 of the Type I CBASS operons was well-tolerated in *E. coli*, with 2 Type I CBASS operons excluded from further analysis due to significant toxicity (Figure 1B). Using classic phages T2, T4, T6, T5, T7 and the phage P1_vir_, we observed that each Type I CBASS operon provided >100-fold defense against at least one phage, with phages T2 and P1_vir_ being particularly susceptible to Type I CBASS defense and phage T4 being especially resistant (Figure 1C). Interestingly, phages T2 and T4 share >98% sequence identity, suggesting that a potentially small number of phage-intrinsic factors may be responsible for differences in CBASS susceptibility (Figure 1C). Together these results reveal a large diversity of functional Type I CBASS operons in biology and establish that minimal two-gene CBASS operons are sufficient to provide robust anti-phage defense in cells.

### A large-scale Type I CBASS phage challenge screen reveals operon- and phage-specific patterns of defense

We next devised a comprehensive screen to map host-virus specificity of Type I CBASS immunity. Leveraging recent construction of the BASEL collection of coliphage isolates^34^, we systematically tested the ability of each Type I CBASS operon to protect *E. coli* against a panel of 65 diverse phages (Figure 2). Our large-scale phage challenge screen of 975 CBASS operon-virus combinations reveals striking patterns of defense against dsDNA phages. We observed that Type I CBASS operons provide robust protection against an exceptionally broad diversity of phages spanning five distinct viral families (*Drexlerviridae, Siphoviridae, Demerecviridae, Myoviridae* [former], and *Autographiviridae*) including examples of both tailed and non-tailed phages and viruses with lytic and temperate replication cycles (Figure 2). CBASS *Ec*CdnD12–4TM, *Ec*CdnF04– SLATT6, and *Vc*CdnG09–Cap5 Type I CBASS operons exhibit potent ≥10,000-fold restriction against phages from across all five viral families tested in our screen (Figure 2; Table S1). Strongly defensive Type I CBASS operons encode diverse CD-NTase enzymes and distinct transmembrane, nuclease, and phospholipase Cap effector proteins demonstrating that multiple CBASS operon architectures can provide broad protection against phage replication *in vivo*.

**Figure 2.**
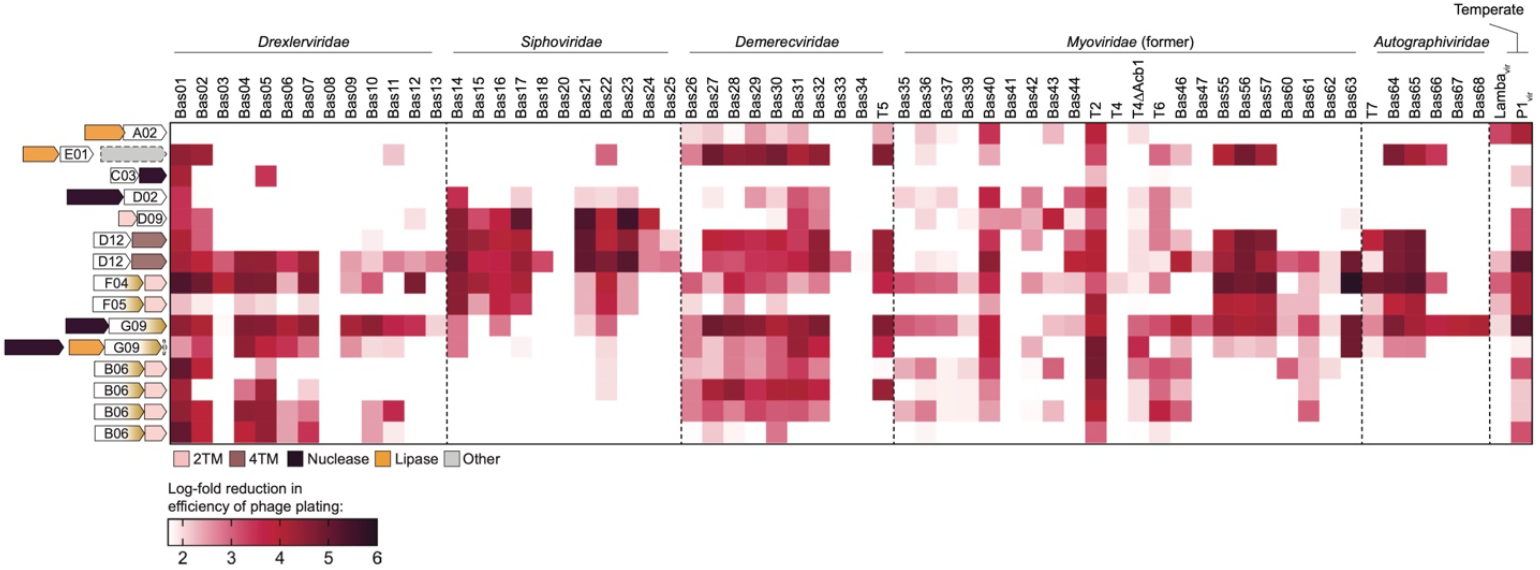
Analysis of large-scale dsDNA phage challenge defines patterns of Type I CBASS defense. Heatmap of log-fold reduction in efficiency of phage plating for each CBASS operon-dsDNA phage combination tested. Expression of Type I CBASS operons was controlled by an arabinose-inducible promoter and expressed using 0.25% (w/v) final concentration for 1 h prior to phage challenge; phage droplets were allowed to dry completely, and plates were incubated overnight at room temperature before imaging and quantification of plaque-forming units (PFUs). A 1.5 log reduction in phage plating threshold was used to define significant phage restriction.

Type I CBASS operons encoding CD-NTases in clades D, F, and G exhibit the broadest protection against dsDNA phage infection and are the only operons which potently restrict the ∼40–60 kb tailed *Siphoviridae* phages and ∼40 kb non-tailed *Autographiviridae* phages (Figure 2). We observed that the defense profiles of clade D, F, G, and *Ec*CdnE01 CD-NTase CBASS operons mirror restriction of the much larger ∼130–140 kb Bas55–57 *Myoviridae* phages (Figure 2), suggesting that these divergent CD-NTases may respond to a conserved molecular cue sensed during infection even among distantly related phages. Comparison of Type I CBASS operons encoding CD-NTase enzymes with an AGS-C domain fusion demonstrates that AGS-C-containing operons are much more broadly protective against T1-like *Drexlerviridae* phages (Figure 2). In contrast, CD-NTases with AGS-C domains exhibit significantly weaker defense against *Siphoviridae* and *Autographiviridae* phages, suggesting that this Ig-like fusion domain can modify the ability of CD-NTases to sense distinct families of phages.

Analysis of phages highly resistant to host immunity suggests that some viruses likely encode immune evasion proteins which counteract Type I CBASS immunity. In four of the five viral families tested, at least one phage exhibits robust resistance to every Type I CBASS operon in our panel. Examples include T1-like phages Bas08 and Bas13, the *Siphoviridae* phages Bas20 and Bas25, T5-like phages Bas33 and Bas34, and *Myoviridae* phages T4 and Bas41 which display broad resistance to Type I CBASS and may express novel anti-CBASS (Acb) immune evasion proteins (Figure 2). In support of these observations, phage T4 is one of the *Myoviridae* phages most resistant to Type I CBASS immunity and is known to encode the anti-defense protein Acb1 which degrades CBASS cyclic nucleotide signals^11^. We observed that Acb1 deletion increases susceptibility of phage T4 to several but not all Type I CBASS operons (Figure 2), in accordance with two recent publications identifying a second anti-CBASS protein Acb2 that functions as a cycle nucleotide sponge to sequester immune signals^14,16^. Intriguingly, phage T2 encodes homologs of both Acb1 and Acb2 anti-CBASS proteins yet is one of the most susceptible phages to Type I CBASS defense (Figure 2), further demonstrating that yet uncharacterized viral genes likely directly or indirectly alter phage susceptibility to CBASS defense.

### Selection for CBASS-resistant phages identifies convergent resistance mutations in structural proteins

We hypothesized that phages may evade Type I CBASS defense by acquiring mutations that alter the proteins or molecular cues that initiate CBASS immune signaling. Using two parallel mutant selection approaches (Figure 3A), we attempted to identify mutant phages that can replicate in the presence of active Type I CBASS (Figure S3A). Agreeing with previous selection attempts to identify CBASS escaper phages in *E. coli*^24^, most phages in our screen failed to spontaneously evolve resistance to Type I CBASS immunity. However, by screening all five families of phages in our large-scale screen against Type I CBASS operons encoding CD-NTases from clades B, D, E, G, and F we ultimately succeeded in isolating 4 unique CBASS-resistant phage species and 6 distinct mutant isolates from the diverse *Drexlerviridae* Bas04, Bas05, Bas11 and Bas13 phages (Figure 3B– D). Remarkably, whole genome sequencing analysis revealed that every CBASS-resistant phage isolate we identified harbored substitution mutations in the scaffolding prohead protease gene required for capsid assembly and maturation (Figure 3B).

**Figure 3.**
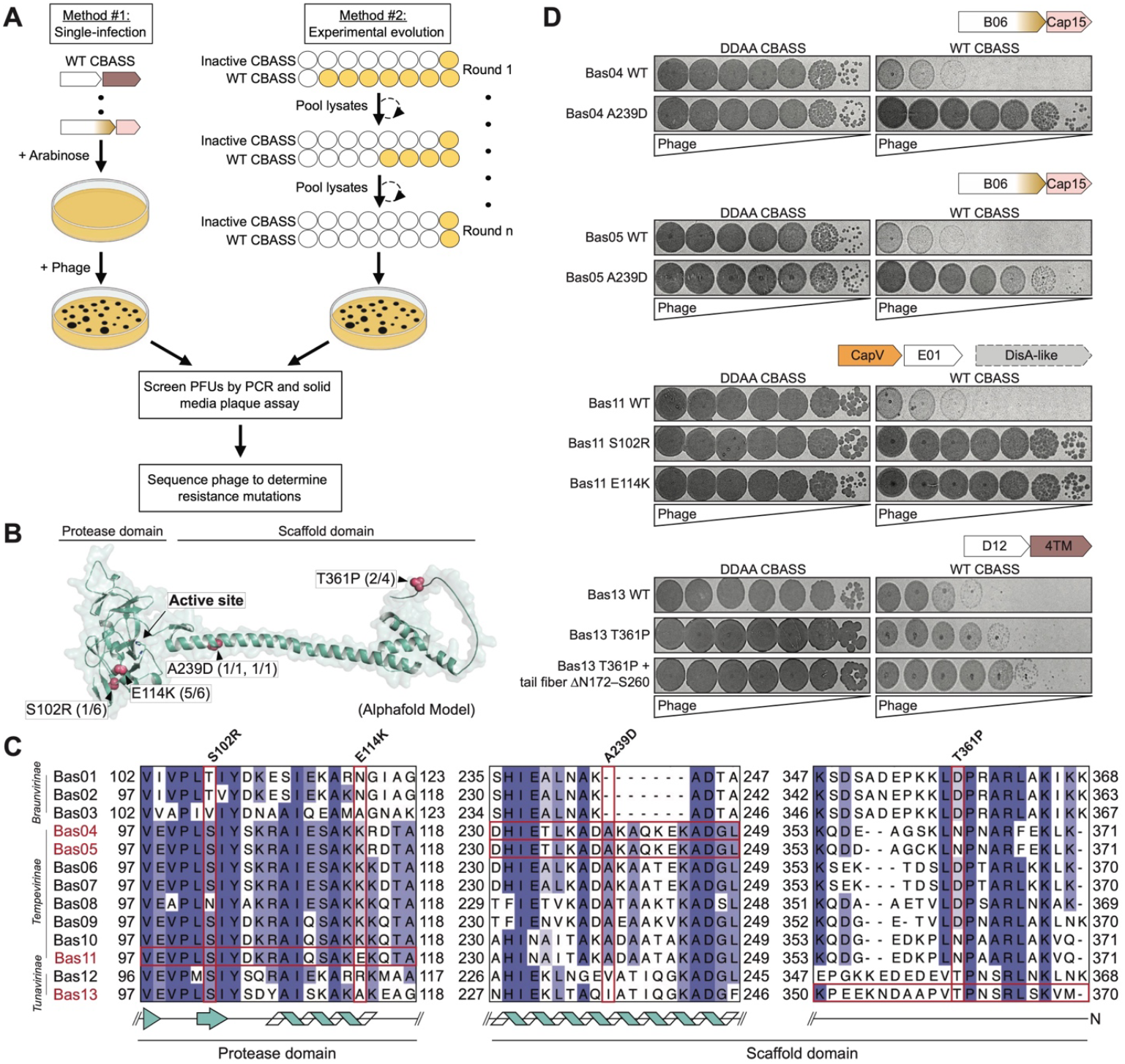
Diverse capsid maturation prohead protease mutations enable evasion of Type I CBASS immunity. (A) Cartoon schematic representation of parallel forward genetic screening methods for CBASS-resistant mutant phage selection in solid (method #1) and liquid media (method #2). (B) Predicted structure of the Drexlerviridae phage prohead protease with CBASS escaper phage mutations identified in this study depicted in red spheres. Numbers represent the number of phage mutants isolated with each mutation. (C) Sequencing analysis of Drexlerviridae phage prohead proteases highlighting substitution mutations by vertical boxes and the phage(s) which harbor each mutation denoted by horizontal boxes. Drexlerviridae subfamily groups are labeled on the Y-axis and cartoon representations below depict predicted protein secondary structure in those regions. (D) Solid media plaque assays showing CBASS escaper phage isolate resistance phenotypes in cells expressing the WT CBASS operon which they were selected against or catalytically inactive CD-NTase CBASS operons which contain two Asp to Ala (DDAA) substitutions in the nucleotidyltransferase active site.

Prohead proteases and capsid scaffolding proteins are highly conserved viral proteins shared across distantly related families of prokaryotic and eukaryotic viruses^35,36^. In *Drexlerviridae* T1-like phages, the prohead protease is comprised of an extended C-terminal alpha-helical scaffolding domain responsible for facilitating capsid head assembly and an N-terminal enzymatic protease domain essential for internal processing of capsid and scaffolding proteins prior to phage dsDNA genome packaging and capsid expansion^36,37^. We mapped escape mutations on a modeled structure of the Bas13 prohead protease and observed that substitutions occur in both protease and scaffolding domains of the phage protein (Figure 3B). Bas11 phage escape mutations cluster in the N-terminal protease domain in a region directly adjacent to the enzyme active site with 5 of 6 isolated Bas11 mutants containing the E114K substitution particularly close to the catalytic residue S121 (Figures 3B and S5C). In contrast, Bas04, Bas05, and Bas13 mutant phages each contain substitutions in the scaffolding domain of the protein (Figure 3B). Bas04 and Bas05 A239D mutations occur near the mid region of the scaffolding domain major alpha-helical spine and the Bas13 T361P substitution maps to the extreme C-terminus of the scaffold domain in an extension predicted to be unstructured (Figure 3B). Sequence analysis demonstrates that most prohead protease escape mutations occur in variable regions of the protein (Figure 3C); interestingly, all other phages in the *Tempevirinae* phage subfamily (Bas04–Bas11) already encode a lysine residue at the same position as the Bas11 E114K escape mutation, suggesting that these phages may have adapted to previous exposure to CBASS or other immune defense (Figure 3C).

We next compared replication of each phage mutant isolate to the parental WT phage in cells expressing the Type I CBASS operon which was used for selection. All Bas04, Bas05, and Bas11 escaper phages exhibit completely restored replication in the presence of functional Type I CBASS operon they were selected against, resulting in a 10,000– 100,000-fold recovery compared to the wildtype virus (Figure 3D). The Bas13 isolate containing only a prohead protease T361P substitution exhibited a more modest ∼10-fold increase in efficiency of plating on WT CBASS-expressing cells (Figure 3D). Notably, several escaper phages including prohead protease mutants Bas04 A239D, Bas11 E114K, Bas11 S102R, and Bas13 T361P contain only a single point mutation, demonstrating that in some cases a single substitution in the prohead protease gene is sufficient to evade Type I CBASS immunity but that acquired resistance varies by mutations (10-fold in the case of the Bas13 T361P single mutant isolate and >10,000-fold in Bas04 and Bas11 resistant isolates) (Figure 3D).

We observed that the Bas05 escaper phage and one Bas13 escaper phage also possessed mutations in one additional protein, but these substitutions were not identified in isolation without prohead protease substitutions, suggesting that the prohead protease mutation preceded these events. The Bas05 CBASS-resistant phage isolate also harbors an N53K mutation in the hypothetical protein ORF78. In a Bas13 mutant isolate containing the prohead protease T361P mutation we identified with an additional ΔN172–S260 deletion mutation in the lateral tail fiber protein (ORF29) that occurs between two 7 bp regions of microhomology within the protein coding region (Figure S3B). Notably, the Bas13 tail-fiber deletion results in an additional 100-fold gain in fitness compared to the isolate harboring the prohead protease T361P mutation alone demonstrating that secondary mutations in structural proteins can further augment resistance to Type I CBASS (Figure 3D). Together our data reveal a novel role for the virion scaffolding maturation prohead protease fusion protein in controlling phage susceptibility to Type I CBASS immunity and demonstrate that mutations in conserved structural proteins are sufficient for phages to evade CBASS immunity without additional anti-defense genes.

### Bas13 prohead protease scaffold domain expression can induce CBASS *Ec*CdnD12-dependent cellular toxicity but does not stimulate CD-NTase activity *in vitro*

We next determined if phage virions or viral structural proteins alone are sufficient to stimulate CD-NTase activity *in vitro*. We were able to express and purify CD-NTase enzymes from the CBASS *Ec*CdnE01–CapV–DisA-like and *Ec*CdnD–4TM operons used in our resistant phage selection experiments. The CD-NTase *Ec*CdnE01 exhibited robust activity *in vitro* in the absence of ligand, but we did not observe stimulation of cyclic nucleotide signal synthesis with either *Ec*CdnE01 or *Ec*CdnD12 in the presence of purified Bas13 virions, prohead protease, or major capsid proteins (Figures 4A and S4C–D). A recent study demonstrated that a Clade E CD-NTase CdnE03 from *Staphylococcus schleiferi* can be stimulated by RNA from phage-infected cells^25^. We therefore additionally tested RNA and DNA purified from Bas11- or Bas13-infected cells but failed to observe an increase in enzymatic activity in either CD-NTase, suggesting that RNA-based enzyme stimulation may not represent a broadly conserved mode of CD-NTase activation (Figure 4A).

**Figure 4.**
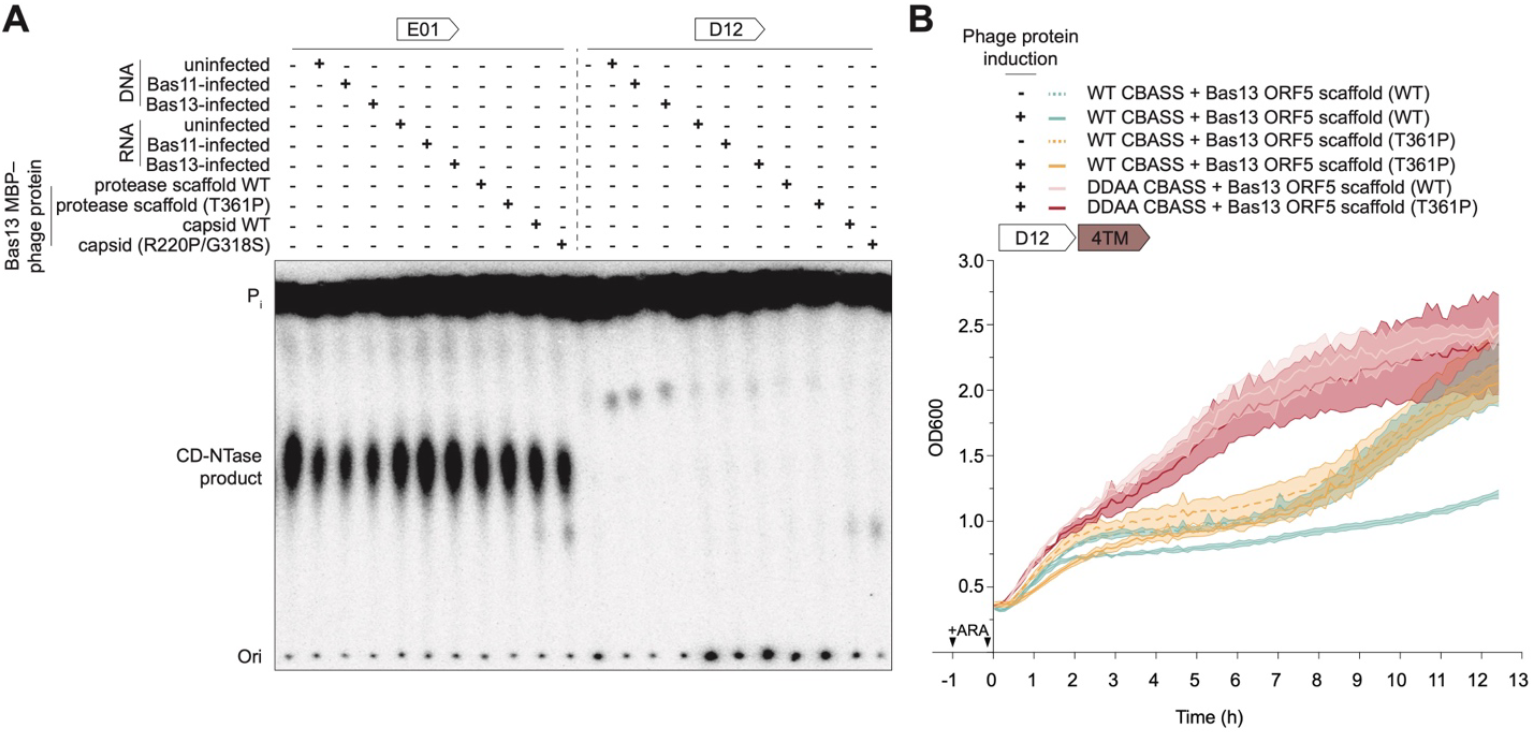
The Bas13 prohead protease scaffold domain induces cellular toxicity in an *Ec*CdnD12 CBASS-dependent manner *in vivo*. (A) Analysis of *Ec*CdnE01 and *Ec*CdnD12 reactions with indicated ligands denoted by a “+” sign. Reactions were labeled with α^32^P-NTPs and terminated by addition of phosphatase to remove terminal phosphates and visualized using thin layer chromatography. Data are representative of at least two biological replicates. (B) *In vivo* CBASS toxicity assay growth curves of WT (green and teal) or inactive DDAA (red and pink) CBASS EcCdnD12–4TM expressing cells in combination with WT or point mutant Bas13 prohead protease scaffold domain (T222–M370). Dotted lines indicate no phage protein IPTG induction. Data are representative of at least three biological replicates and error bars displayed as shaded regions are the standard deviation of three technical replicates.

Phage structural proteins oligomerize during virion assembly^36,37^, and we hypothesized that our *in vitro* assays with monomeric proteins may not recapitulate complex interactions that occur *in vivo* during viral replication. We therefore next tested whether expression of Bas13 structural proteins was sufficient to induce activation of the CBASS *Ec*CdnD12–4TM *in vivo*. Remarkably, we observed that expression of the Bas13 prohead protease scaffold domain (T222–M370) was sufficient to induce CBASS-mediated cellular toxicity (Figure 4B). In contrast, no CBASS-mediated cellular toxicity was observed upon expression of the Bas13 major capsid protein (Figure S4F). Co-expression of the Bas13 prohead protease scaffold domain in cells expressing a CBASS operon encoding a mutated CD-NTase at two catalytic aspartate residues (DDAA) resulted in no toxicity, confirming that the prohead protease scaffold domain itself is not toxic to cells and that the effect we see is dependent on CD-NTase activity (Figure 4B, solid red and pink lines). As an additional control, we withheld the IPTG inducer driving phage protein expression and observed that expression of the CBASS *Ec*CdnD12–4TM operon alone delayed culture growth, likely due to higher than native protein levels when expressed from a heterologous plasmid system, but these conditions were significantly less toxic compared to co-expression with the Bas13 prohead protease scaffold over the same period (Figure 4B, dotted teal line), demonstrating that the cellular toxicity phenotype is also dependent on phage protein expression. Finally, we observed that introduction of the Bas13 prohead protease T361P *de novo* CBASS resistance mutation was sufficient to abrogate *Ec*CdnD12 CBASS-mediated cellular toxicity (Figure 4B). Together, these results suggest that some Type I CBASS operons may directly sense the phage proteins involved in late-stage viral assembly but that CD-NTase activation may require recognition of complex events beyond interaction of monomeric proteins.

### Phage resistance to Type I CBASS immunity is specific to operons with sequence or architectural similarity

To begin to characterize mechanisms of CBASS evasion, we next analyzed if phage mutations provide general or operon-specific resistance to Type I CBASS immunity. We challenged each phage mutant isolate against every Type I CBASS operon in our panel and observed that phage resistance to Type I CBASS immunity is highly operon-specific (Figures 5A and S5A). Phage Bas04 and Bas05 isolates with prohead protease scaffold domain mutations evade the *Psp*CdnB06– Cap15 CBASS operon they were selected against but remain susceptible to *Xp*CdnC03– NucC, *Ec*CdnD12–4TM, *Ec*CdnF04–SLATT6, *Ec*CdnF05–SLATT6, *Vc*CdnG09–Cap5, and *Vv*CdnG09–CapV–Cap5 Type I CBASS-mediated restriction (Figure 5A). Strikingly, we observed that phage Bas04 and Bas05 mutants also acquire 1,000–10,000-fold resistance to other CBASS operons encoding closely related clade B06 CD-NTase enzymes *Ea*CdnB06, *Asp*CdnB06 and *Pp*CdnB06. The Clade B06 CD-NTase enzymes in these four CBASS operons are ∼50–75% similar at the amino acid level, suggesting a common mechanism of phage resistance to these CBASS operons with high sequence similarity (Figures 1A and 5A). Similarly, all phage Bas11 prohead protease enzymatic domain mutant isolates resist the CBASS *Ec*CdnE01 operon they were selected against by ≥100-fold but are unable to fully overcome *Pp*CdnB06-, *Ec*CdnD12-, *Vc*CdnG09, and *Vv*CdnG09-mediated CBASS immunity (Figure 5A). These results are indicative of a general pattern in which mutant phages evade CBASS systems closely related to the system they were selected against but fail to overcome more distantly related Type I CBASS operons encoding a distinct CD-NTase enzyme (Figure 5A).

**Figure 5.**
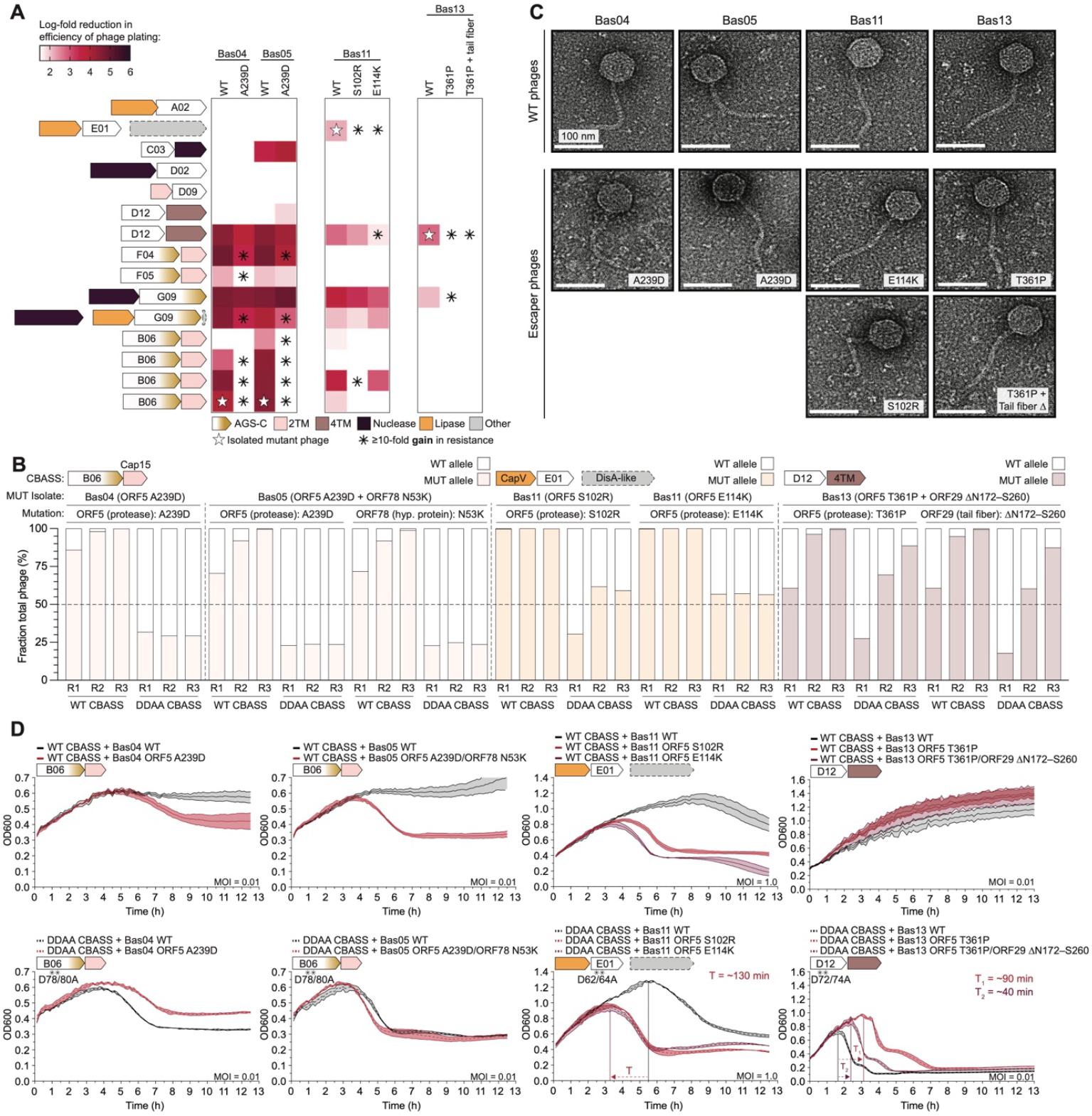
Phage resistance to Type I CBASS immunity is operon-specific and has diverse effects on viral fitness. (A) CBASS escaper phage cross-resistance analysis heatmap of log-fold reduction in efficiency of phage plating of mutant phages against screened Type I CBASS operons. Asterisks indicate where ≥10-fold resistance was acquired by a mutant phage isolate against the indicated CBASS operon. (B) Sequencing analysis of WT and mutant phage competition assays. WT and indicated mutant phages were combined in equal ratios and used to infect WT CBASS or inactive DDAA CBASS-expressing cultures followed by passaging for 3 rounds as indicated on the X-axis. The Y-axis represents the fraction of total phage harboring the mutation indicated above based on DNA sequencing reads. (C) Liquid media phage challenge assays for WT (black lines) and escaper phages (red, maroon lines) in cells expressing WT (solid lines) or inactive CD-NTase (DDAA) CBASS (dotted lines) at the indicated multiplicity of infection (MOI). Horizontal lines with arrows highlight significant changes between WT/mutant phage lysis timing. Representative experiments of at least two biological replicates are shown and error bars for technical triplicates are shown as the shaded region surrounding each line. (D) Negative stain electron micrographs of purified WT or mutant phage virions high lighting indistinguishable capsid head structures.

Other specific examples of CBASS resistance in our analysis reveal that phage escaper mutations can affect susceptibility to Type I CBASS that share operon architectural similarity. We observed that phage Bas04 and Bas05 mutant isolates acquire ≥10-fold resistance to some additional Type I CBASS operons that encode clade F04/05 and clade G09 CD-NTase enzymes which have high sequence divergence compared to the clade B06 CBASS operon they were selected against (Figures 5A and S5A). Interestingly, these divergent CBASS operons share the presence of an AGS-C Ig-like fusion domain, further suggesting that this domain may play a conserved role in phage sensing or regulating cyclase activity. Similar results were observed with phage Bas11 and Bas13 escaper mutants acquiring cross-resistance against some Type I CBASS operons distantly related to the system they were selected against. Bas11 S102R and E114K escaper mutant isolates selected against the CBASS *Ec*CdnE01 operon exhibit ≥100-fold resistance to *Pp*CdnB06–Cap15 and ≥10-fold resistance against *Ec*CdnD12–4TM, respectively (Figure 5A). Likewise, although WT phage Bas13 is only susceptible to two Type I CBASS operons in our screen, we observed that Bas13 escaper phages selected against *Ec*CdnD12 CBASS additionally gained ∼10-fold resistance against the divergent *Vc*CdnG09–Cap5 CBASS system (Figure 5A). Notably, no Bas11 or Bas13 CBASS escaper mutant isolates exhibit a broadly increased susceptibility to highly dissimilar Type I CBASS operons apart from select instances (Figure 5A). Together, these results reveal that CBASS escaper mutations can provide resistance to more distantly linked operons that display related operon architecture and, in rarer cases, may confer cross-resistance to highly divergent CBASS operons with no evident sequence or architectural similarity.

### Phage resistance to CBASS immunity is not associated with a significant cost to viral fitness

To determine the impact of CBASS escaper mutant phages on phage fitness, we next co-infected cultures with WT and mutant phage isolates and used next-generation sequencing to monitor viral replication over three rounds of infection. As expected, in the presence of functional Type I CBASS immunity, most escaper phages rapidly outcompeted parental WT strains and reached w>99% of viral reads by the end of round 3 of replication (Figures 5B and S5). In contrast, co-infection experiments performed in CBASS deficient cells revealed that in the absence of immune pressure CBASS escaper mutations have a variable impact on viral fitness. Bas04 and Bas05 escaper phage isolates with a prohead protease scaffold domain A239D mutation exhibited neutral or slightly reduced replication fitness compared to WT isolates (Figure 5B). Likewise, Bas11 phages with prohead protease enzymatic domain S102R and E114K mutations replicated to approximately the same level as WT phages in CBASS deficient cells demonstrating that these CBASS escaper mutations do not strongly affect viral fitness (Figure 5B). Finally, the Bas13 CBASS escaper phage isolate with a prohead protease T361P mutation and the lateral tail fiber protein ΔN172–S260 deletion outcompeted WT Bas13 co-infection cultures with or without functional Type I CBASS defense demonstrating that this isolate has a general advantage compared to the WT parental phage (Figure 5B). Overall, these results reveal that CBASS resistance mutations are largely innocuous to general viral fitness and represent a specific adaptation to exposure to CBASS immunity.

We also observed that some phages can acquire additional structural protein mutations under sustained Type I CBASS immune pressure. During passage, a proportion of Bas11 S102R escaper phages acquired additional prohead protease substitutions including the A59D and Q116R mutations (Figure S5B,C). The Q116R prohead protease mutation is predicted to affect local electrostatic interactions similar to the E114K mutation observed in other Bas11 isolates but the A59D mutation is not near the active site (Figure S5C) and the potential effects of this mutation on protein function are less clear. Interestingly, during co-infection with WT Bas13 and the Bas13 prohead protease T361P single mutant escaper isolate in CBASS *Ec*CdnD12-expressing cells, we found that the Bas13 T361P mutation was lost during the first round of infection, indicating that this escaper mutation is insufficient to outcompete WT phage under co-culture conditions (Figure S5D). However, our sequencing analysis detected novel substitutions in the phage major capsid protein (ORF8) by the third round of in infection where we observed a ∼50% frequency of the major capsid protein G318S substitution in the otherwise WT phage population following loss of the prohead protease substitution. To confirm these findings, we isolated clonal Bas13 phages from round 3 of the co-infection mixed population and identified two new Bas13 CBASS escaper mutant isolates with a major capsid protein N273D single mutation and R220P/G318S double mutation which both separately restored efficiency of plating by ≥10,000-fold in WT CBASS-expressing cells (Figure S5E). Notably, major capsid mutations have previously been identified as escaper mutations that enable the *Pseudomonas* phage PaMx41 to evade Type II CBASS immunity^14^. These results demonstrate that mutations in multiple structural proteins allow phages to evade Type I CBASS defense and further suggest that escaper mutations affect a late stage of virion assembly to enable host immune evasion.

### CBASS resistance mutations can affect phage infection kinetics

The identification of CBASS resistance mutations in three essential proteins required for virion assembly^36,37^ led us to ask whether mutant phage isolates were affected in virion morphology. We analyzed purified phage virions using negative-stain electron microscopy and observed that Bas04, Bas05, Bas11, and Bas13 prohead protease and Bas13 prohead protease / lateral tail fiber double-mutant CBASS escaper phages produced virions that are morphologically indistinct from their WT counterparts at this resolution (Figure 5C)^34^. We were unable to visualize intact Bas13 capsid mutant phages by negative-strain EM, suggesting that these CBASS escaper mutations may lead to the formation of destabilized virions. Supporting this possibility, the Bas13 capsid R220P, N273D, and G318S mutations are predicted to occur at the capsid hexameric interface as predicted in the *Pseudomonas* phage PaMx41 described above and also previously observed with mutations in the *E. coli* phage T5 major capsid protein demonstrated to evade cyclic mononucleotide-based Pycsar immunity but at significant fitness cost to the virus (Figure S5F)^14,33^. These results demonstrate that some CBASS escaper mutations may impact phage virion assembly but that most phage isolates which fully evade Type I CBASS immunity are unaffected in infectious virion morphology.

To further define how CBASS resistance mutations in structural proteins may impact late stages of viral infection we next measured infection kinetics in liquid media in cells expressing the WT CBASS operon used for selection or an operon which has been inactivated by two aspartate to alanine substitutions in the CD-NTase nucleotidyltransferase core (DDAA CBASS). At a low multiplicity of infection, we observed that in the absence of Type I CBASS immunity Bas04, Bas05, Bas11, and Bas13 WT phages lyse *E. coli* cultures and induce culture collapse by ∼6 h of infection (Figure 5D). As expected, cells expressing functional Type I CBASS are resistant to Bas04, Bas05, Bas11, and Bas13 WT phage infection and sustain culture growth compared to inactive DDAA CBASS-expressing cultures (Figure 5D). Bas04 and Bas05 isolates with prohead protease scaffold domain CBASS resistance mutations fully lyse *E. coli* cultures with or without functional Type I CBASS immunity by ∼6 h post infection and exhibit unchanged replication dynamics compared to WT phages. However, Bas11 S102R and E114K isolates with prohead protease enzymatic domain mutations lyse cultures ∼130 min faster than WT phage Bas11, revealing that these mutations lead to faster infection kinetics in a CBASS-independent manner (Figure 5D). Further supporting that CBASS escaper mutations can impact phage infection dynamics, we also observed altered culture collapse timing in Bas13 mutant isolates. The Bas13 prohead protease T361P single mutant isolate which is only modestly resistant to CBASS EcCdnD14– 4TM immunity collapses *E. coli* cultures ∼90 min later than WT phage Bas13 (Figures 3D and 5D). Interestingly, the Bas13 CBASS escaper double mutant isolate with the prohead protease T361P substitution and tail fiber protein ΔN172–S260 deletion exhibits faster infection kinetics than the prohead protease mutant alone with culture collapse timing intermediately restored to Bas13 WT levels (Figure 5D). Finally, we also characterized the Bas13 capsid mutant isolates identified by our sequencing analyses and find that these mutant phages lyse WT CBASS-expressing cultures by ∼6 h, but both Bas13 capsid N273D single mutant and R220P/G318S double mutant isolates exhibit a ∼60 min delay in culture lysis compared to the WT phage Bas13 without functional CBASS immune pressure. Taken together, our results reveal that likely multiple paths exist to altering the rate of late-stage virion assembly to enable phages to evade CBASS immunity and provide detailed characterization of escaper phages which demonstrates that CBASS resistance mutations are a specific adaptation to CBASS immunity and can dramatically alter culture lysis kinetics.

## DISCUSSION

Our comprehensive analysis of Type I CBASS provides a new foundation to explain CBASS immune specificity and viral evasion. Previous experiments have analyzed subsets of CBASS operons and focused on Type II and Type III systems with additional auxiliary regulatory genes^1,5,9,14–16^. We present nearly 1,000 Type I CBASS operon-phage pairings and demonstrate that diverse, minimal CBASS operons potently restrict replication of a broad range of dsDNA phages (Figure 1, Figures 2 and S2). Furthermore, our characterization of 15 new Type I CBASS operons creates a resource of functional Type I CBASS systems that span major clades A–G of CD-NTase enzymes and eight diverse Cap effector protein classes.

Comprehensive analysis of Type I CBASS operon phage restriction profiles reveals specific patterns of defense against dsDNA phages and new correlates between phage restriction and CBASS operon architecture. Some Type I CBASS operons exhibit an exceptionally broad range of defense against diverse dsDNA phages, particularly operons from *Escherichia coli* SLK172 and *Escherichia coli* BIDMC103 encoding CD-NTase enzymes from clades D and F, respectively. Notably, no *in vitro* cyclic nucleotide synthesis activity has previously been observed for any clade F CD-NTase enzyme^2^, but our results now demonstrate that these enzymes are functional in cells and can restrict replication of diverse dsDNA phages (Figures 1 and 2). We also found that restriction of some phages by Type I CBASS is highly CD-NTase clade-specific, suggesting that some CD-NTase enzymes may respond to molecular cues that are conserved in distinct subsets of phages (Figures 1 and 2). Interestingly, Type I CBASS operons which encode a CD-NTase enzyme fused to an AGS-C Ig-like domain more broadly restrict replication of phages from the *Drexlerviridae* family (Figure 2). AGS-C domains are additionally found fused to cyclic CMP and cyclic UMP cyclases in Pycsar immunity where they also correlate with clade-specific distribution among PycC enzymes^33^, suggesting that this domain may have a conserved role in controlling aspects of phage specificity in both CBASS and Pycsar bacterial immunity. Finally, our analysis demonstrates that some phages including Bas08, Bas20, Bas34, T4, and Bas68 are exceptionally resistant to Type I CBASS defense. In each case, Type I CBASS operons potently restrict closely related family members, indicating that these resistant phages may encode anti-CBASS proteins that inhibit host immune defense (Figure 2). In support of this prediction, phage T4 becomes markedly more susceptible to Type I CBASS immunity upon deletion of the known anti-CBASS nuclease Acb1 (Figure 2)^11^. These results, along with the recent discovery of Acb2 as second family of anti-CBASS proteins^14,16^, suggest that novel inhibitors of CBASS immunity remain to be discovered.

Our results reveal a convergence of mutations in viral structural proteins that enable phages to escape Type I CBASS immunity. Surprisingly, 6 out of 8 unique phage mutant isolates from our selection experiments harbored substitution mutations in the prohead protease gene required for virion capsid assembly and maturation (Figure 3). We show that co-expression of the CBASS *Ec*CdnD12– 4TM operon with the phage Bas13 prohead protease scaffolding domain is sufficient to induce cellular toxicity in a CBASS-dependent manner (Figure 4). CBASS-dependent toxicity is relieved by a single scaffold domain T361P substitution, demonstrating that *de novo* substitutions that give rise to immune resistance do not require additional phage-encoded anti-defense genes to evade Type I CBASS defense. Notably, a recent preprint reported that *Staphylococcus* phage Φ80a_vir_ can evade Type I CBASS *Ssc*CdnE03–Cap15 immunity via mutations in gp46—a viral capsid scaffold protein homologous to the *Drexlerviridae* scaffolding domain identified in this study (Figure S4F)^25^. Phage Φ80a_vir_ gp46 escaper mutations map to a region distinct from the prohead protease scaffold or protease domain substitutions identified in our study, further highlighting a convergence of CBASS escaper mutations across highly divergent phage species (Figure S4A). In our study we additionally identify major capsid and tail-fiber mutations that allow phages to evade Type I CBASS, however, expression of the Bas13 major capsid protein alone was not sufficient to induce cellular toxicity in cells expressing a functional CBASS *Ec*CdnD12–4TM operon or stimulate activity of two CD-NTases *in vitro* (Figures 4A and S4F). This is consistent with previous reports in CBASS and Pycsar defense systems demonstrating that major capsid protein mutations enable immune evasion but are insufficient to initiate immune signaling *in vivo*^14,33^. Finally, we discovered that some mutations in the phage prohead protease—but not others—lead to changes in phage culture lysis timing, implicating multiple evasion strategies via mutagenesis of a single phage protein. Together, these results demonstrate that mutations in diverse phage structural proteins can alter the late-stages of virion assembly to enable resistance to CBASS defense.

Our results identify the phage prohead protease as a novel determinant of CBASS activation and evasion. These results suggest that some CD-NTases may directly recognize phage structural proteins to initiate CBASS immune signaling. However, the scaffold domain was not sufficient to stimulate CD-NTase activation in vitro (Figure 4), indicating that if *Ec*CdnD12 or other CD-NTases directly interact with the prohead protease that binding may require recognition of an oligomeric or intermediate conformation that exists during phage virion assembly. Defining the basis of these potential interactions and the mechanism of CD-NTase activation is now an important goal for future analysis. Additionally, it remains to be fully understood if acquiring mutations in the prohead protease and scaffolding proteins represents a broadly applicable mode of CBASS evasion. These results advance our understanding of initiation and escape of bacterial CBASS immunity and provide a foundation to discover novel CBASS activating cues and phage-encoded inhibitors.

## METHODS

### Bacterial strains and growth conditions

All bacterial strains used are of the *Escherichia coli* species. *E. coli* TOP10 cells were used for plasmid cloning and maintenance and were routinely grown at 37°C in liquid and on solid media supplemented with appropriate antibiotic for plasmid maintenance (100 mg mL^−1^ ampicillin and 34 mg mL^−1^ chloramphenicol). Cloning of toxic phage genes was conducted in media supplemented with 1% glucose to suppress inducible expression constructs and increase cell viability. All solid and liquid media phages challenge assays were conducted in *E. coli* BW25113 cells, a kind gift from T. Bernhardt (Harvard Medical School).

### Type I CBASS CD-NTase phylogenetic analysis and operon selection

Available CD-NTase sequences were acquired from Millman et al^8^. Sequences were trimmed using GeniusPrime software to include the N-terminal-most 300 amino acid residues; iTOL was used to align sequences and generate a rooted phylogenetic tree followed by addition of clade information from Whitley et al^2^. Bioinformatically annotated AGS-C domains were identified using data from Millman et al^8^ and incorporated into the phylogenetic tree as black circles; bacterial CD-NTase clade information is depicted as inner shaded region. We selected Type I CBASS operons exclusively form gammaproteobacterial species for use in *E. coli* phage challenge experiments and genes adjacent to core CBASS genes were included if they could not be ruled out as a function of genomic distance^38^. CBASS operons were synthesized (Genescript) and cloned into a custom pBAD vector.

### Solid media phage challenge small-droplet plaque assays

Competent bacterial cells were transformed with indicated plasmids using standard transformation protocols. Single colonies or glycerol stocks were used to inoculate LB cultures supplemented with 5 mM MgCl_2_, 5 mM CaCl_2_, 0.1 mM MnCl_2_, and appropriate antibiotic. Liquid cultures were grown for 4–6 h to OD_600_ = 1.0–1.5 and then normalized to OD_600_ = 0.3 in 0.8% top agar (5 mL for small plates and 15 mL for larger plates) containing 0.25% (w/v) arabinose and left for 1 h at room temperature to set top agar and induce CBASS expression. 10-fold serial dilutions of stock phage were prepared in SM Buffer (100 mM NaCl, 50 mM Tris-HCl pH 7.5, and 8 mM MgSO_4_) and 3 μL droplet were plated on the surface of the bacterial top agar lawn; droplets were allowed to dry completely (10–20 min), and plates were left to incubate overnight at room temperature. Plates were imaged the following day using a BioRadTM Chemidoc and quantification of efficiency of plating was determined by counting plaque-forming units (PFUs) compared to inactive CBASS control.

### Single-infection selection for CBASS-resistant phage mutants in solid media

Competent *E. coli* BW25113 cells were transformed with indicated CBASS operon-expressing plasmids using standard transformation protocols. Single colonies or glycerol stocks were used to inoculate LB cultures supplemented with ampicillin used at 100 μg mL^−1^. Cultures were incubated for 4–6 h at 37°C shaking until reaching an OD_600_ of 1.0–1.5 and then back-diluted to OD_600_ = 0.3–0.5 in 0.8% low-melting point molten top agar supplemented with 0.25% (w/v) arabinose, 5 mM MgCl_2_, 5 mM CaCl_2_, 0.1 mM MnCl_2_, and 100 μg mL^−1^ ampicillin (final concentrations). High titer phage stocks were then added to the above mixture at final dilutions between 10^−6^ and 10^−10^ depending on lysate titer to achieve discernable single plaque-forming units (PFUs). Final mixtures were briefly vortexed to ensure a homogenous mixture, immediately plated on standard LB agar plates and incubated overnight at room temperature (∼25°C). The following day, individual PFUs were isolated and mixed in 50 μL SM Buffer, vortexed briefly and spun down, then used as input for phage genotyping by PCR and verification of CBASS resistance by small-droplet solid media plaque assay. Phage clones positive for both tests were propagated by inoculating 1.5 mL log phage *E. coli* BW25113 cultures (OD_600_ = 0.3–0.5) transformed with the WT CBASS operon used for selection whose expression was pre-induced using 0.25% arabinose (^w^/_v_, final conc.) ∼30 min prior to phage infection and incubated at 37°C overnight followed by pelleting cell debris and 0.22 μm filter clarification. Clarified phage lysates were checked again for resistance to the CBASS that was used for selection by solid media plaque assay before preparation for sequencing. 1 mL of clarified phage lysates were treated with DNase I and RNase A for 90 min at 37°C without agitation to remove free nucleic acids. Nucleases were inactivated by addition of 20 mM (final concentration) EDTA, and samples flash-frozen in LN_2_ and shipped overnight on dry ice to SeqCenter who performed phage genomic DNA extractions, sequencing library preparations, and high-throughput Illumina DNA sequencing (2 × 151 bp paired end reads).

### Experimental evolution selection for CBASS-resistant phage mutants in liquid media

This method was adapted from Srikant et al, 2022^39^. Competent *E. coli* BW25113 cells were transformed with indicated WT CBASS or inactive CBASS operon-expressing plasmids using standard transformation protocols. Single colonies or glycerol stocks were used to inoculate LB cultures supplemented with ampicillin. Cultures were incubated for 2–3 h at 37°C shaking to establish culture growth and followed by addition of 0.25% (w/v) (final conc.) for 1 h. Culture OD_600_ was measured and 10^5^ colony-forming units (CFUs) WT or inactive CBASS-expressing cells were added to every other row of a 96-well plate in a 200 μL volume of LB media supplemented with 0.25% (w/v) arabinose, 5 mM MgCl_2_, 5 mM CaCl_2_, 0.1 mM MnCl_2_ (final conc.). The first two rows are both seeded with inactive CBASS-expressing cells as a control. For the first round of infection, the entire plate is infected with the same serially diluted stock phage at a high MOI of 10^2^ in 10 μL volumes of SM buffer or with buffer alone for one column which receives no phage. 96-well plates were sealed with a breathable film and cells were incubated at 28°C shaking overnight and harvested the followed day. For each pair of rows (1 pair of rows of inactive CBASS only and 5 pairs of rows with WT and inactive CBASS), up to 5 cleared wells from each row starting from the lowest MOI are pooled from each pair of rows to generate a unique mixed lysate population. For subsequent rounds of infection, 96-well plates are seeded as above, and each population lysate (control or 1–5) was used to infect the same population of each subsequent plate. Individual plaque-forming units were isolated and verified by PCR and plaque assay prior to preparation and shipment for whole genome sequencing as described above.

### Whole genome sequencing bioinformatic analysis

For phage whole genome sequencing analysis, paired-end reads were pre-processed using Cutadapt^40^ to trim adapter readthrough and filter poor-quality reads. Reads were mapped to the parental phage genome using bwa (aln, sampe)^41^ and variant single-nucleotide polymorphisms were called using BCFtools (mpileup, call, filter)^42^. Per-position coverage was obtained using SAMtools^43^ depth and coverage maps were plotted with a rolling 150-nt window size average of per-nucleotide coverage.

### Recombinant protein expression and purification

cGAS/DncV-like nucleotidyltransferases (CD-NTases) and phage Bas13 proteins were purified from *E. coli* as previously described^2,9,18^. Briefly, genes were cloned into custom pET expression vectors containing N-terminal 6×His-SUMO2 or 6×His-MBP tags, respectively, by Gibson assembly using HiFi DNA Assembly Master Mix (NEB). Expression plasmids were transformed into BL21 (DE3) RIL cells (Agilent) and plated onto MDG media (1.5% Bacto agar, 0.5% glucose, 25 mM Na_2_HPO_4_, 25 mM KH_2_PO_4_, 50 mM NH_4_Cl, 5 mM Na_2_SO_4_, 0.25% aspartic acid, 2–50 μM trace metals, 100 μg mL^−1^ ampicillin, 34 μg mL^−1^ chloramphenicol). After overnight incubation at 37°C, three colonies were used to inoculate a 30 mL MDG culture overnight (12–16 h) at 37°C, shaking). 1L M9ZB cultures (47.8 mM Na_2_HPO_4_, 22 mM KH_2_PO_4_, 18.7 mM NH_4_Cl, 85.6 mM NaCl, 1% casamino acids, 0.5% glycerol, 2 mM MgSO_4_, 2–50 μM trace metals, 100 μg mL^−1^ ampicillin, 34 μg mL^−1^ chloramphenicol) were then inoculated with the MDG starter culture at OD_600_ of 0.05 and grown at 37°C, shaking to an OD_600_ of 1.5–3.0 before induction with 0.5 mM isopropyl-β-d-thiogalactoside (IPTG) overnight (12–16 h) shaking. After overnight M9ZB expression, cell pellets were harvested by centrifugation and then resuspended and lysed by sonication in 50 mL lysis buffer (20 mM HEPES-KOH pH 7.5, 400 mM NaCl, 10% glycerol, 30 mM imidazole, 1 mM TCEP). Lysates were clarified by centrifugation at 50,000 × g for 30 min, supernatant was poured over 8 mL Ni-NTA resin (Qiagen), resin was washed with 35 mL lysis buffer supplemented with 1 M NaCl, and protein was eluted with 10 mL lysis buffer supplemented with 300 mM imidazole. Samples were then dialyzed overnight in dialysis tubing with a 14 kDa molecular weight cutoff (Ward’s Science). Proteins were used for biochemical assays and dialyzed in dialysis buffer supplemented with 10% glycerol. Purified proteins were concentrated to >15 mg ml^−1^ using 30-kDa MWCO centrifugal filter units (Millipore Sigma), aliquoted, flash frozen in liquid nitrogen and stored at −80 °C.

### CD-NTase basal activity and activation thin-layer chromatography assays

Thin-layer chromatography was used to analyze cyclic nucleotide second messenger products as previously described^2,11^. 1 μL of purified CD-NTase protein was incubated with 50 μM of each unlabeled nucleotides ATP, CTP, GTP, UTP and 0.5 μL α-^32^P-labeled NTPs (approximately 0.4 μCi each of nucleotide) in cGLR reaction buffer (50 mM Tris-HCl pH 7.5, 100 mM KCl, 10 mM MgCl_2_, and 1 mM 724 TCEP) at 37°C overnight for basal activity tests or for 2 h for testing nucleic acid and protein ligands. Human cGAS incubated with ISD45 was used as a positive control^44^. Reactions were terminated with addition of 0.5 μL of Quick CIP (New England Biolabs) to remove terminal phosphate groups from unreacted nucleotides. Each reaction was analyzed using TLC by spotting 0.5 μL on a 20 cm × 729 20 cm PEI-cellulose TLC plate (Millipore). The TLC plates were developed in 1.5 M KH_2_PO_4_ pH 3.8 until buffer was 1–3 cm from the top of plate and air-dried at room temperature and exposed to a phosphor-screen before imaging with a Typhoon Trio Variable Mode Imager (GE Healthcare).

### *In vivo* CBASS cellular toxicity growth curve plate reader assays

Competent *E. coli* BW25113 cells were co-transformed with WT CBASS operons or CBASS operon which has been inactivated by two CD-NTase catalytic aspartate to alanine substitutions (DDAA) CD-NTase and indicated WT or mutated phage protein-expressing constructs using standard transformation protocols onto LB agar plates containing ampicillin and chloramphenicol. Single colonies were used to inoculate 3 mL LB liquid cultures supplemented with antibiotic. Cultures were incubated for 2–3 h at 37°C shaking to establish culture growth and followed by addition of 0.25% (w/v) (final conc.) for 1 h. OD_600_ measurements were taken for all cultures and a normalized 1.37 × 10^8^ cells was aliquoted for ± phage protein induction conditions. Cells were pelleted by centrifugation at 3,500 × g for 5 min followed by removal of media and washing with 750 μL PBS. Cells were again pelleted by centrifugation and resuspended in 650 μL LB media supplemented with 5 mM MgCl_2_, 5 mM CaCl_2_, 0.1 mM MnCl_2_, 0.125% arabinose and either containing 1 mM IPTG or no IPTG. Each sample is then divided into 3 wells of a 96-well plate in technical triplicate and loaded in the plate reader machine and OD_600_ absorbance was measured in ∼7.5 min increments with intermittent shaking. Data were visualized using GraphPad Prism 9 software; representative images of at least two biological replicates are shown and standard error of technical triplicates is represented as the shaded area around data points.

### WT and mutant phage competition assays in liquid media

Competent *E. coli* BW25113 cells were transformed with indicated WT CBASS operons or operons which harbor two aspartate to alanine substitutions in the CD-NTase active site (DDAA) using standard transformation protocols. Single colonies or glycerol stocks were used to inoculate LB cultures supplemented with 5 mM MgCl_2_, 5 mM CaCl_2_, 0.1 mM MnCl_2_ and appropriate antibiotic(s) (ampicillin used at 100 mg mL^−1^ and chloramphenicol used at 34 mg mL^−1^). Cultures were incubated for 3–4 h at 37°C shaking to establish growth followed by addition of arabinose at 0.25% (final w/v) to induce CBASS expression; cultures were then further incubated for an addition 1 h until early log phage (OD_600_ = 0.3–0.5) and normalized to 4.20 × 10^8^ total CFU in 1.5 mL final volume LB media supplemented as above with trace metals and arabinose. Equal MOI of WT and indicated mutant phages was added to appropriate cultures followed by brief mixing via gentle vortex and incubation overnight (12–16 h). The following day, phage populations were harvested from collapsed cultures by removal of bacterial cell debris via centrifugation for 15 min at 4,000 rpm and lysate clarification via 0.22 μm filtration. The resulting mixed population phage stocks were tittered by small-droplet plaque assay on bacterial lawns of inactive CBASS-expressing cells and used to infect cultures for the next round at the total MOI (WT + mutant phages) of the previous round.

### Negative stain electron microscopy of phage virions

3.5 μL aliquots of 0.22 μm-filtered high-titer phage lysate samples were diluted as necessary in SM buffer and applied onto glow-discharged (30 sec, 30 mA) 400-mesh copper grids (EMS400-Cu) coated with a continuous layer of approximately 5 nm carbon. Samples were allowed to adsorb for 1 minute and then blotted using filter paper (VMR, 28310-081). The grid was then immediately stained with 1.5% uranyl formate solution (Electron Microscopy Science, 22451) and then blotted. The staining procedure was repeated twice with a 1 min incubation with uranyl formate before the final blotting step. The grid was air-dried at room temperature prior to imaging. Electron microscopy images were collected using a FEI Tecnai T12 transmission electron microscope operating at 120 kV and equipped with a Gatan UltraScan 895 (4k x 4k) CCD camera at a nominal magnification of 42,000× corresponding to a calibrated pixel size of 2.61 Å and a defocus value of −3.5 μm.

### Liquid media phage challenge bacterial growth curve plate reader assays

Competent bacterial cells were transformed with indicated plasmids using standard transformation protocols. Single colonies or glycerol stocks were used to inoculate LB cultures supplemented with 5 mM MgCl_2_, 5 mM CaCl_2_, 0.1 mM MnCl_2_ and appropriate antibiotics (ampicillin used at 100 mg mL^−1^ and chloramphenicol used at 34 mg mL^−1^). Liquid cultures were grown to log-phase OD_600_ = 0.3–0.5 and then normalized to OD_600_ = 0.3 and aliquoted in 200 μL volumes of supplemented LB media in technical triplicate in a 96-well plate format. Triplicate cultures were then inoculated with diluted stock phage in 10 μL volumes at the indicated multiplicity of infection (MOI). The 96-well plate was then sealed with a liquid-impermeable film and loaded in the plate reader machine and OD600 absorbance was measured in ∼7.5 min increments with intermittent shaking. Data was manipulated using Microsoft Excel and visualized using GraphPad Prism 9 software; representative images of at least two biological replicates are shown and standard error of technical triplicates is represented as the shaded area around data points.

### Quantification and Statistical Analysis

Details of quantification and statistical analysis are listed in the Figure legends. Experiments were performed with at least 3 independent replicates. Statistical analyses were conducted using GraphPad Prism Version 9.1. Data are plotted with error bars representing standard error of the mean (SEM).

## Author Contributions

Experiments were designed by D.R.-B. and P.J.K with input from S.J.H. D.R.-B. performed phylogenetic analysis and cloning of Type I CBASS operons. S.J.H. propagated bacteriophage strains for large-scale screening with assistance from D.R.-B. D.R.-B. performed all solid and liquid media phage challenge assays, mutant phage selection experiments, phage fitness comparison studies, and *in vivo* CBASS toxicity growth curves. J.M.G. analyzed all whole genome sequencing data with support from A.S.Y.L. H.T. performed purifications of recombinant proteins and *in vitro* CD-NTase basal activity tests. D.R.-B. and H.T. performed *in vitro* thin-layer chromatography CD-NTase activation experiments. J.G. conducted negative stain-EM structural studies with samples prepared by D.R.-B. The manuscript was written by D.R.-B. and P.J.K., and all authors contributed to editing the manuscript and support its conclusions.

## Acknowledgements

The authors are grateful to members of the Kranzusch lab for helpful comments and discussion. The authors also thank A. Harms (University of Basel) and M. Baum (Harvard Medical School) for sharing bacteriophage samples, S. Srikant (Massachusetts Institute of Technology) for helpful discussions on mutant selection, and L. DuBois (Harvard Medical School) for help optimizing two-plasmid expression systems. The work was funded by grants to P.J.K. from the Pew Biomedical Scholars program, the Burroughs Wellcome Fund PATH program, The Mathers Foundation, The Mark Foundation for Cancer Research, the Parker Institute for Cancer Immunotherapy, and the National Institutes of Health (1DP2GM146250), grants to A.S.Y.L. from the Pew Biomedical Scholars Program, The Mathers Foundation, the V Foundation, and the National Institutes of Health (1R35GM142527), and grants to S.S. from the National Institutes of Health (1DP2GM137415 and 1R01AG073277). D.R.-B. is supported by the National Science Foundation (NSF) Graduate Research Fellowship Program (GRFP), S.J.H. is supported through a Cancer Research Institute Irvington Postdoctoral Fellowship (CRI3996), and J.M.G. is supported by a Ford Foundation Predoctoral Fellowship.

---

**Table S1.**
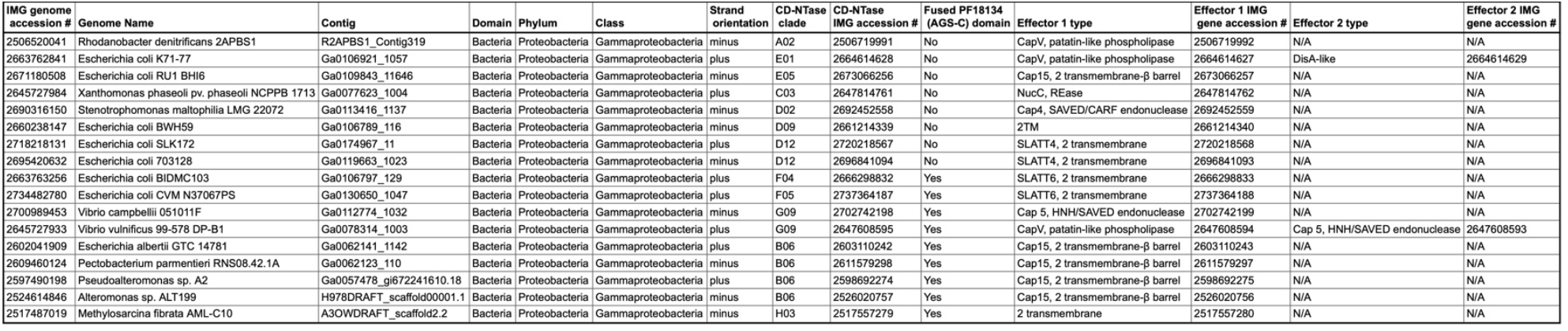
related to Figure 1.

**Figure S2.**
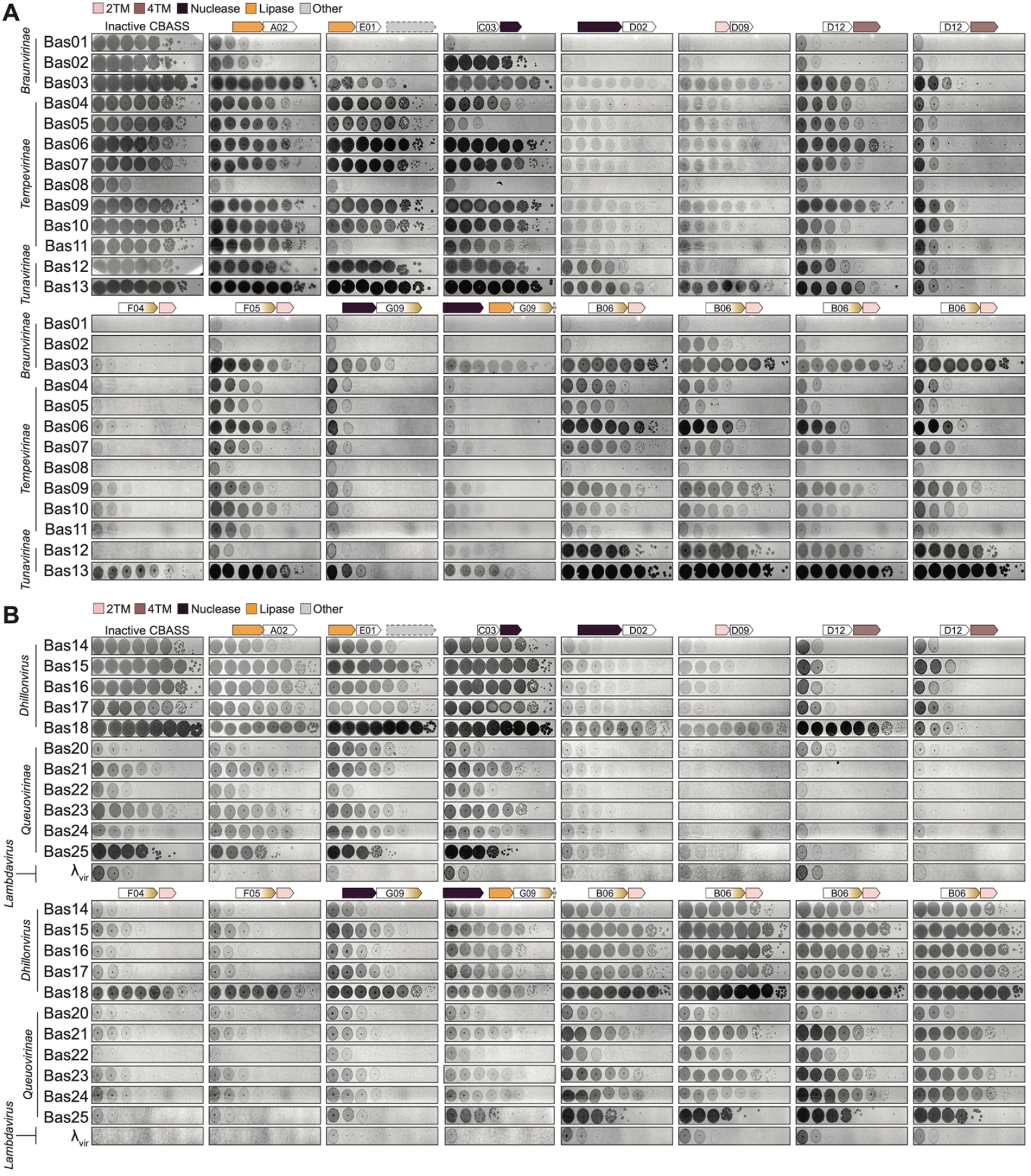
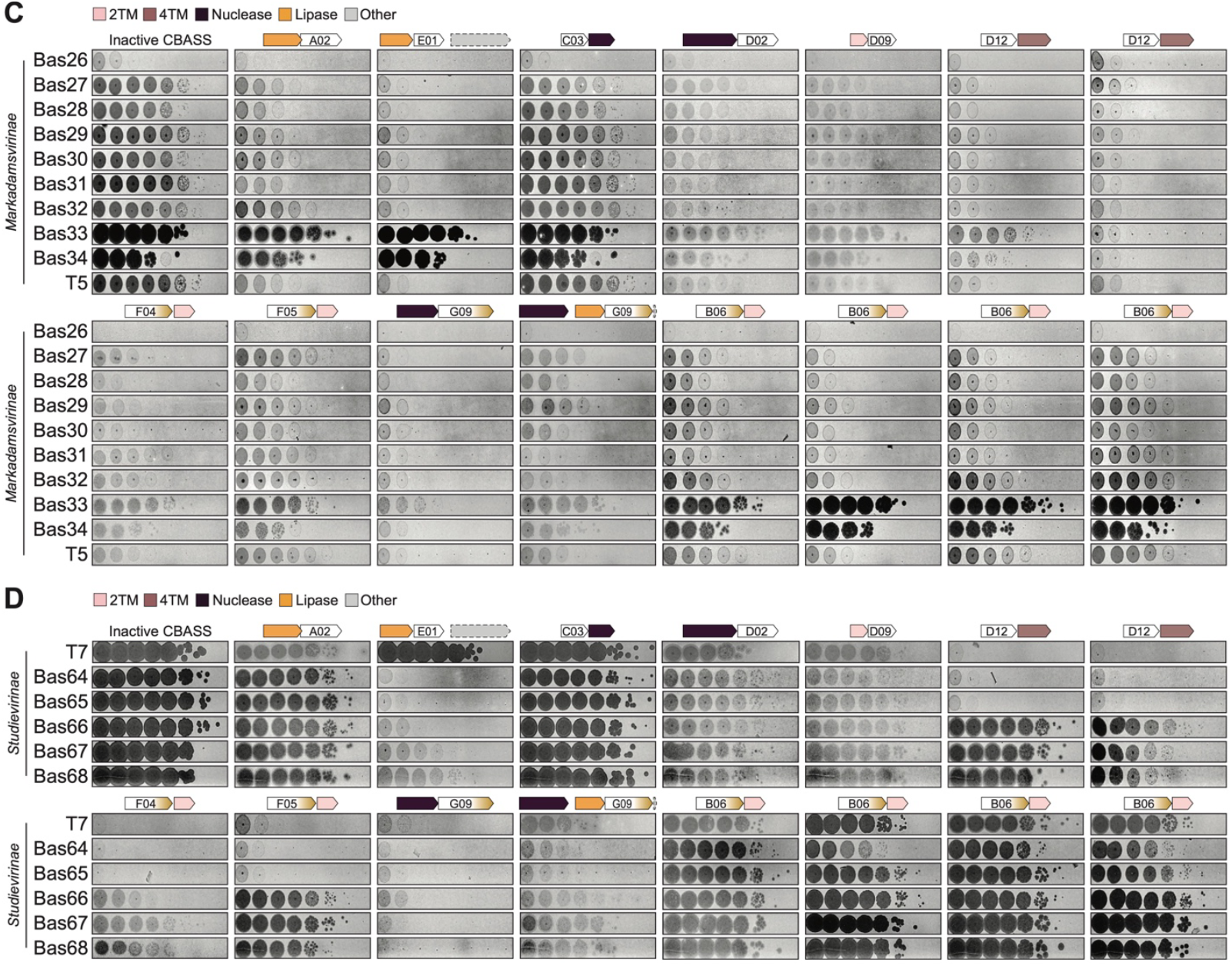
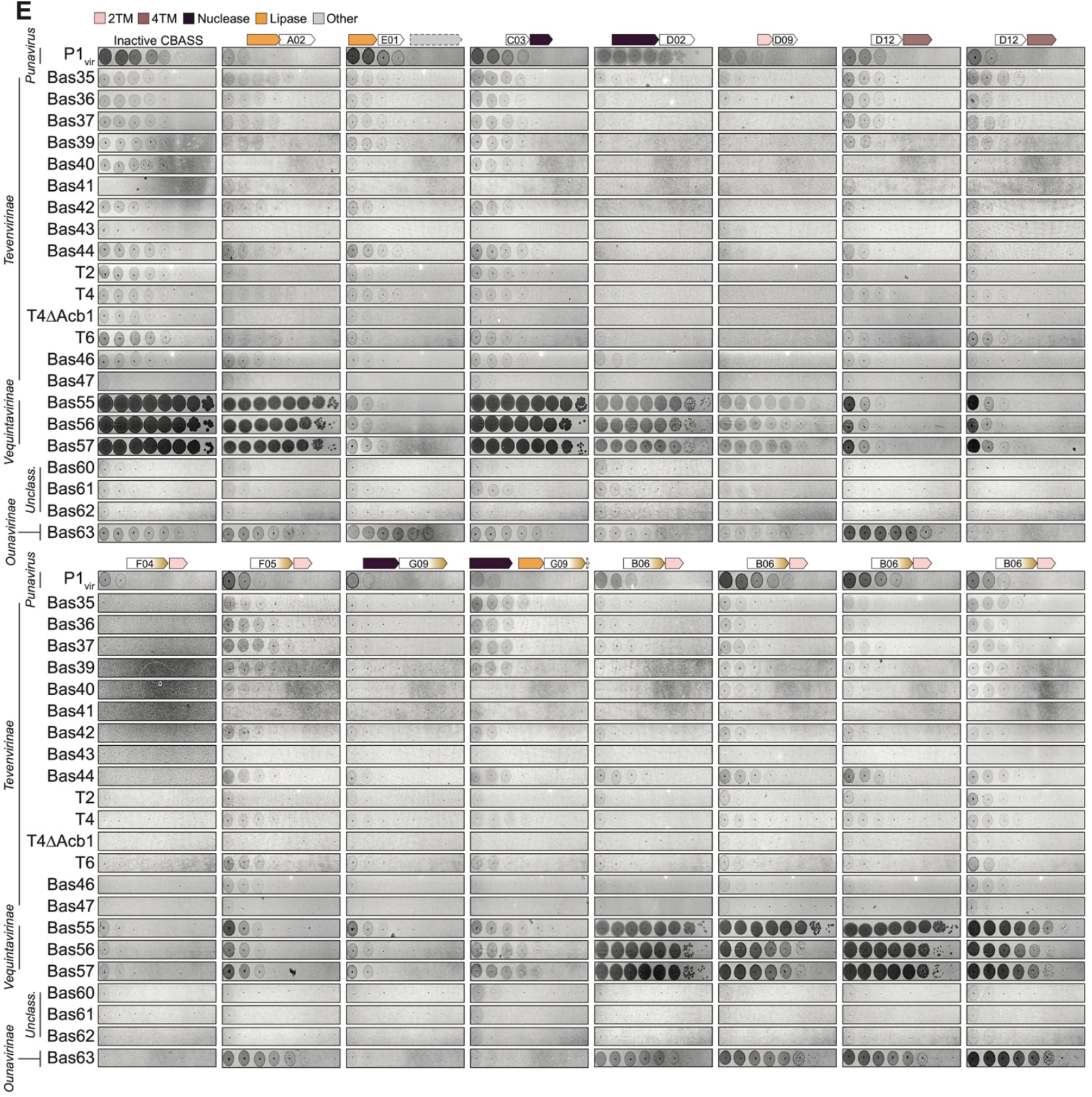
Type I CBASS operons encoding diverse CD-NTase enzymes and cap effector proteins broadly and potently restrict dsDNA phage infection in a heterologous expression system, related to Figure 2. (A–E) Primary solid media small droplet plaque assay data for *in vivo* screening of Type I CBASS operons used in this study. CBASS operons (X-axis) are controlled by an arabinose inducible promoter and expressed using 0.25% (w/v, final) arabinose for 1 h prior to phage challenge with droplets of serially diluted phage indicated on the Y-axis. Phage droplets are allowed to dry completely and incubated overnight (12–16 h) at room temperature (∼25°C) followed by imaging. Plaque-forming units (PFUs) are enumerated and log-fold reduction in efficiency of phage plating (EOP) is determined compared to cells expressing the previous validated effector-null inactive CBASS control^10^. Viral subfamilies are labeled on the Y axis. Heatmap in Figure 2 is the average log-fold reduction in efficiency of phage plating of at least two biological replicates.

**Figure S3.**
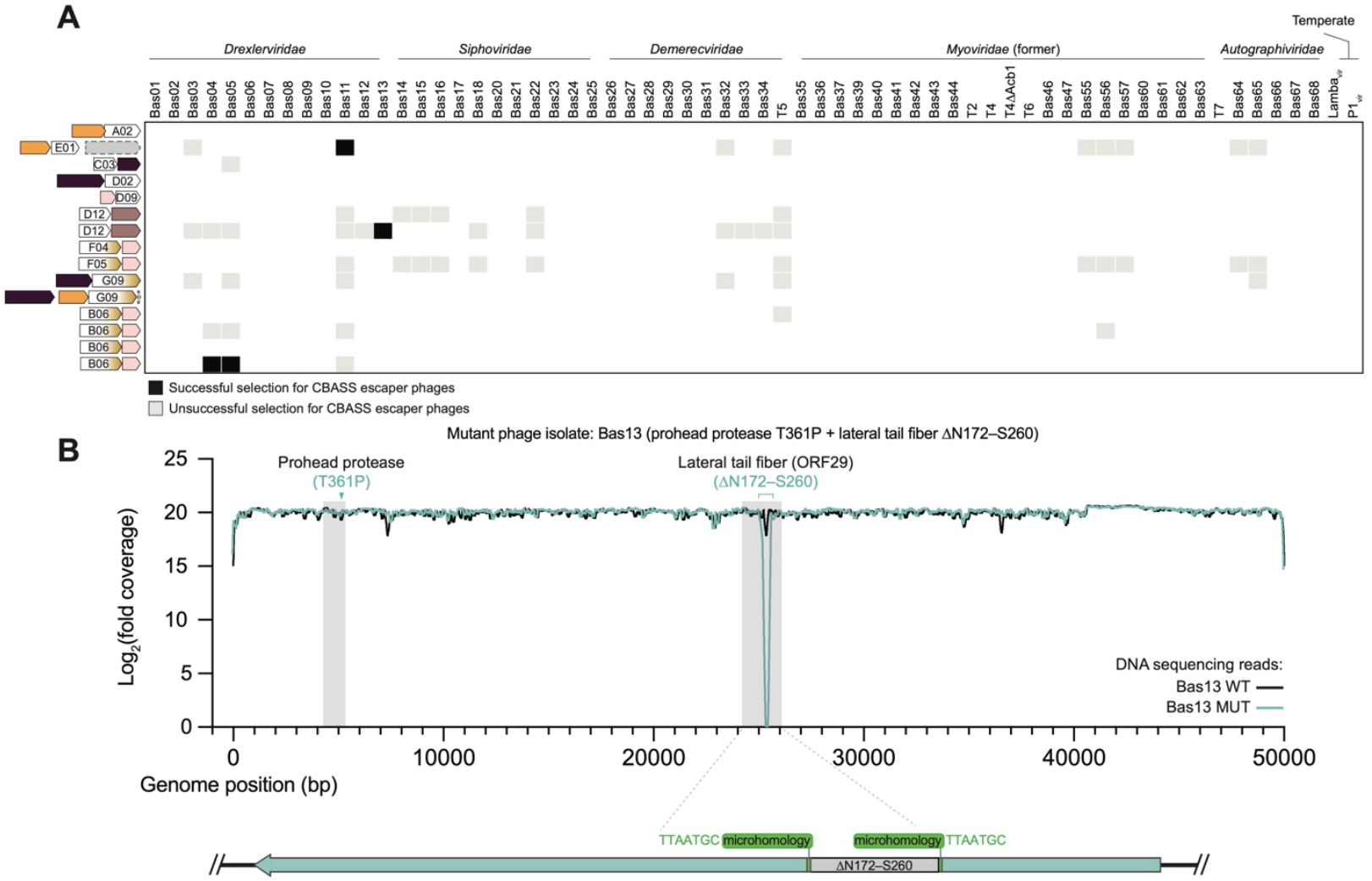
CBASS-resistant phage selections reveal diverse Drexlerviridae escaper phage mutations, related to Figure 3. (A) Grid of CBASS operon-phage pairings tested in this study highlighting successful (black boxes) and attempted (gray boxes resistant phage selections. (B) Whole genome Log_2_ sequencing read coverage of the Bas13 ORF5 (prohead protease) T361P / ORF29 (lateral tail fiber) ΔN172– S260 double mutant isolate highlighting microhomology-mediated deletion mutation.

**Figure S4.**
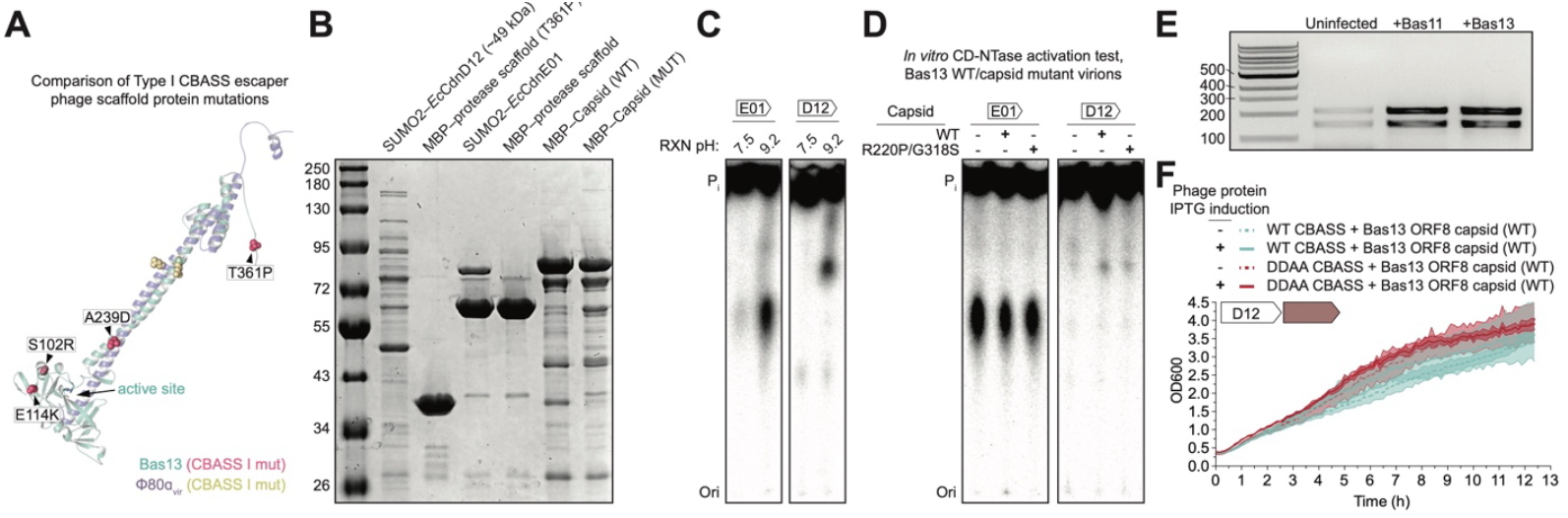
*In vitro* and *in vivo* analysis of the ability of phage structural proteins and purified virions to stimulate *Ec*CdnD12 and *Ec*CdnE01 CD-NTase activity, related to Figure 4. (A) Structural prediction analysis comparing Bas13 ORF5 prohead protease and ϕ80α_vir_ gp46 scaffold protein with escaper mutations identified in each study displayed in as spheres. (B) SDS-PAGE and Coomassie Blue stain analysis of purified CD-NTase and phage structural protein. (C) Basal CD-NTase activity test at pH 7.5 and 9.2. Reactions were incubated overnight with all four NTPs and trace α^32^P-labeled nucleotides and analyzed by thin-layer chromatography. (D) CBASS *Ec*CdnD12 and *Ec*CdnE01 *in vitro* activation assay with WT and mutant phage Bas13 virions showing no change in CD-NTase activity with addition of ligand. (E) Total RNA extracted using Trizol reagent from uninfected, Bas11-infected, and Bas13-infected separated on an 0.8% TAE agarose gel. (F) CBASS *Ec*CdnD12–4TM cellular toxicity assay. Co-expression of WT *Ec*CdnD12 CBASS with the Bas13 major capsid protein does not lead to CBASS-dependent cellular toxicity.

**Figure S5.**
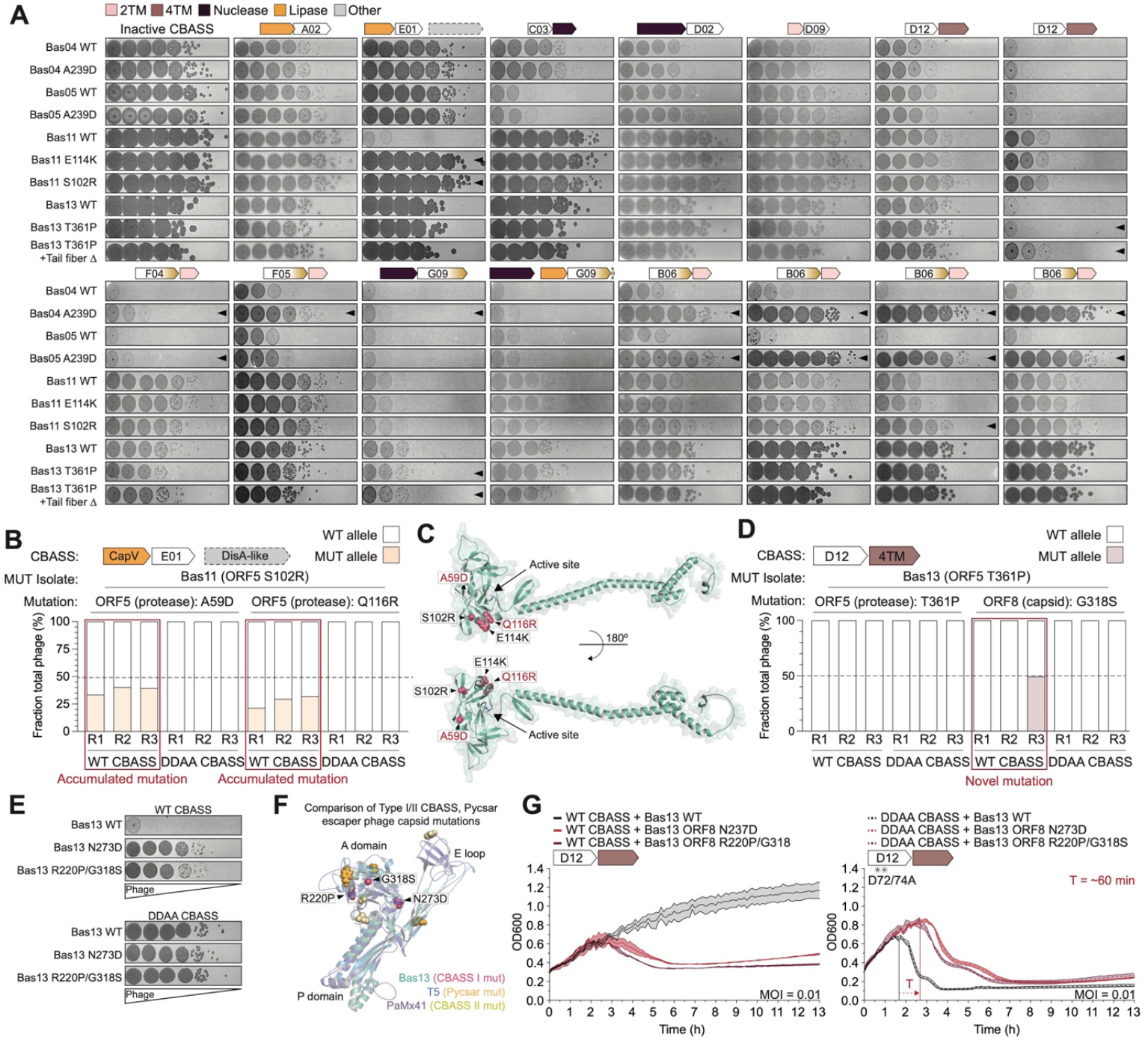
Acquired cross-resistance to Type I CBASS is CD-NTase clade-specific and reveals accumulated and novel mutations, related to Figure 5. (A) Small droplet plaque assay data for CBASS escaper phage cross-resistance analysis to determine log-fold reduction in efficiency of phage plating of mutant phages against all screened Type I CBASS operons. Arrows indicate where ≥10-fold resistance was acquired by a mutant phage isolate against the indicated Type I CBASS operon. (B) Additional sequencing analysis of WT and Bas11 ORF5 prohead protease S102R mutant isolate competition assays highlighting accumulation of two novel A59D and Q116R substitutions under sustained WT CBASS immune pressure. WT mutant phages were combined in equal ratios and used to infect WT CBASS- or DDAA (inactive) CBASS-expressing cultures followed by passaging for 3 rounds as indicated on the X-axis. The Y-axis represents the fraction of total phage harboring the mutation indicated above based on DNA sequencing reads. (C) Predicted structure of the Drexlerviridae phage Bas11 prohead protease enzymatic domain CBASS resistance mutations depicted as red spheres. Originally identified escaper mutations are in black text and mutations accumulated under sustained CBASS immune pressure are highlighted in red text. (D) Sequencing analysis of WT and Bas13 ORF5 prohead protease T361P mutant isolate competition assays showing loss of the prohead protease T361P mutation after round 1 of infection. Appearance of a novel substitution in the major capsid protein (ORF8) is highlighted in a red box. (E)Solid media plaque assays showing identified Bas13 capsid mutant CBASS escaper phage resistance phenotypes in cells expressing the WT CBASS EcCdnD12–4TM operon which they were selected against or a CBASS operon which contains two aspartate to alanine (DDAA) substitutions in the CD-NTase active site. (F) Comparison of the predicted structures of major capsid proteins from Bas13, Pseudomonas phage PaMx41^14^, and E. coli phage T5^33^. Mutations identified in this study are labeled and shown as spheres with structure and mutations color combinations depicted in the key in the bottom right. The three common domains of the HK97 major capsid fold are labeled without boxes. (G) Liquid media phage challenge assays for WT (black lines) and escaper phages (red, maroon lines) in cells expressing WT (solid lines) or catalytically inactive CD-NTase (DDAA) CBASS (dotted lines) at the indicated multiplicity of infection (MOI). Horizontal lines with arrows highlight significant changes between WT/mutant phage lysis timing. Representative experiments of at least two biological replicates are shown and error bars for technical triplicates are shown as the shaded region surrounding each line.

